# FMO1 catalyzes the production of taurine from hypotaurine

**DOI:** 10.1101/750273

**Authors:** Sunil Veeravalli, Ian R. Phillips, Rafael T. Freire, Dorsa Varshavi, Jeremy R. Everett, Elizabeth A. Shephard

**Affiliations:** Department of Structural and Molecular Biology, University College London, London WC1E 6BT, UK; School of Biological and Chemical Sciences, Queen Mary University of London, London E1 4NS, UK; Medway Metabonomics Research Group, University of Greenwich, Chatham Maritime, Kent ME4 4TB, UK

## Abstract

Taurine is one of the most abundant amino acids in mammalian tissues. It is obtained from the diet and by *de novo* synthesis, from cysteic acid or hypotaurine. Despite the discovery in 1954 that the oxygenation of hypotaurine produces taurine, the identification of an enzyme catalyzing this reaction has remained elusive. In large part this is due to the incorrect assignment, in 1962, of the enzyme as a NAD-dependent hypotaurine dehydrogenase. For more than 55 years the literature has continued to refer to this enzyme as such. Here we show, both *in vivo* and *in vitro*, that the enzyme that oxygenates hypotaurine to produce taurine is flavin-containing monooxygenase 1 (FMO1). Metabolite analysis of the urine of *Fmo1*-null mice by ^1^H NMR spectroscopy revealed a build-up of hypotaurine and a deficit of taurine in comparison with the concentrations of these compounds in the urine of wild-type mice. *In vitro* assays confirmed that FMO1 of human catalyzes the conversion of hypotaurine to taurine utilizing either NADPH or NADH as co-factor. FMO1 has a wide substrate range and is best known as a xenobiotic- or drug-metabolizing enzyme. The identification that the endogenous molecule hypotaurine is a substrate for the FMO1-catalyzed production of taurine resolves a long-standing mystery. This finding should help establish the role FMO1 plays in a range of biological processes in which taurine or its deficiency is implicated, including conjugation of bile acids, neurotransmitter, anti-oxidant and anti-inflammatory functions, the pathogenesis of obesity and skeletal muscle disorders.

## Introduction

Taurine (2-aminoethanesulfonic acid) is one of the most abundant amino acids in mammalian tissues (1). It is an organic osmolyte involved in cell volume regulation (1) and has a variety of cytoprotective and developmental roles, particularly in neurological and ocular tissues (2). It is also involved in the formation of bile salts (1) and the modulation of intracellular calcium concentration (3). Taurine is considered an essential substance in mammals and its deficiency has been implicated in several pathological conditions; however, the mechanism by which the amino acid exerts its effects is not understood.

Taurine is obtained from the diet and, by *de novo* synthesis, from cysteic acid (4) or hypotaurine (5). Hypotaurine is itself an organic osmolyte and cytoprotective agent (6) and acts as an anti-oxidant to scavenge highly reactive hydroxyl radicals (7). The oxygenation of hypotaurine to produce taurine was discovered in 1954 (5). Later the enzyme that converts hypotaurine to taurine was reported to be an NAD-dependent hypotaurine dehydrogenase (8), but was not isolated or characterized. Subsequently, Oja *et al.* (9) noted that the production of taurine in tissue extracts was optimal at pH 9.0 and was stimulated by oxygenation. These authors concluded that the enzyme that converts hypotaurine to taurine was not an NAD-dependent hypotaurine dehydrogenase. By overlooking the work of Oja *et al.* (9), and giving credence to the earlier study (8), the enzyme that catalyzes the conversion of hypotaurine to taurine has continued to be reported as a hypotaurine dehydrogenase, utilising NAD as cofactor (EC 1.8.1.3). Because an enzyme that catalyzes the reaction has not been identified or isolated, the conversion of hypotaurine to taurine has sometimes been referred to as non-enzymatic.

Here we show, both *in vivo*, through the use of a knockout-mouse line, and *in vitro*, by assays of human enzymes, that oxygenation of hypotaurine to produce taurine is catalyzed by flavin-containing monooxygenase 1 (FMO1).

## Results

Previous phenotypic analysis (10–13) has identified metabolic differences between mice in which the *Fmo1, Fmo2* and *Fmo4* genes had been deleted (*Fmo1*^*−/−*^, *2*^*−/−*^, *4*^*−/−*^ mice) (10) and wild-type animals. As an extension of this work we have used one-dimensional (1D) ^1^H NMR spectroscopy to compare the urinary metabolic profiles of the knockout mouse line and wild-type animals. Analysis of the urine of male and female *Fmo1*^*−/−*^, *2*^*−/−*^, *4*^*−/−*^ mice revealed signals at 2.66 and 3.37 ppm, corresponding to those of hypotaurine. Such signals were markedly lower in the urine of wild-type mice (Fig. 1A, B). Signals at 3.28 and 3.43 ppm, corresponding to those of taurine, were detected in the urine of both wild-type and *Fmo1*^*−/−*^, *2*^*−/−*^, *4*^*−/−*^ mice, but their intensities were lower in the latter (Fig. 1A, B). The identities of taurine and hypotaurine in urine samples were confirmed by two-dimensional (2D) NMR (Supporting Information Fig. S1). The urinary ratio of taurine to hypotaurine + taurine was significantly less in *Fmo1*^*−/−*^, *2*^*−/−*^, *4*^*−/−*^ mice than in wild-type mice (*P* <0.0001) (Fig. 1C). The build-up of hypotaurine and the concomitant decrease of taurine in the urine of *Fmo1*^*−/−*^, *2*^*−/−*^, *4*^*−/−*^ mice (Fig. 1A, B, C) suggests that the formation of taurine from hypotaurine is catalyzed by an FMO.

**Figure 1.**
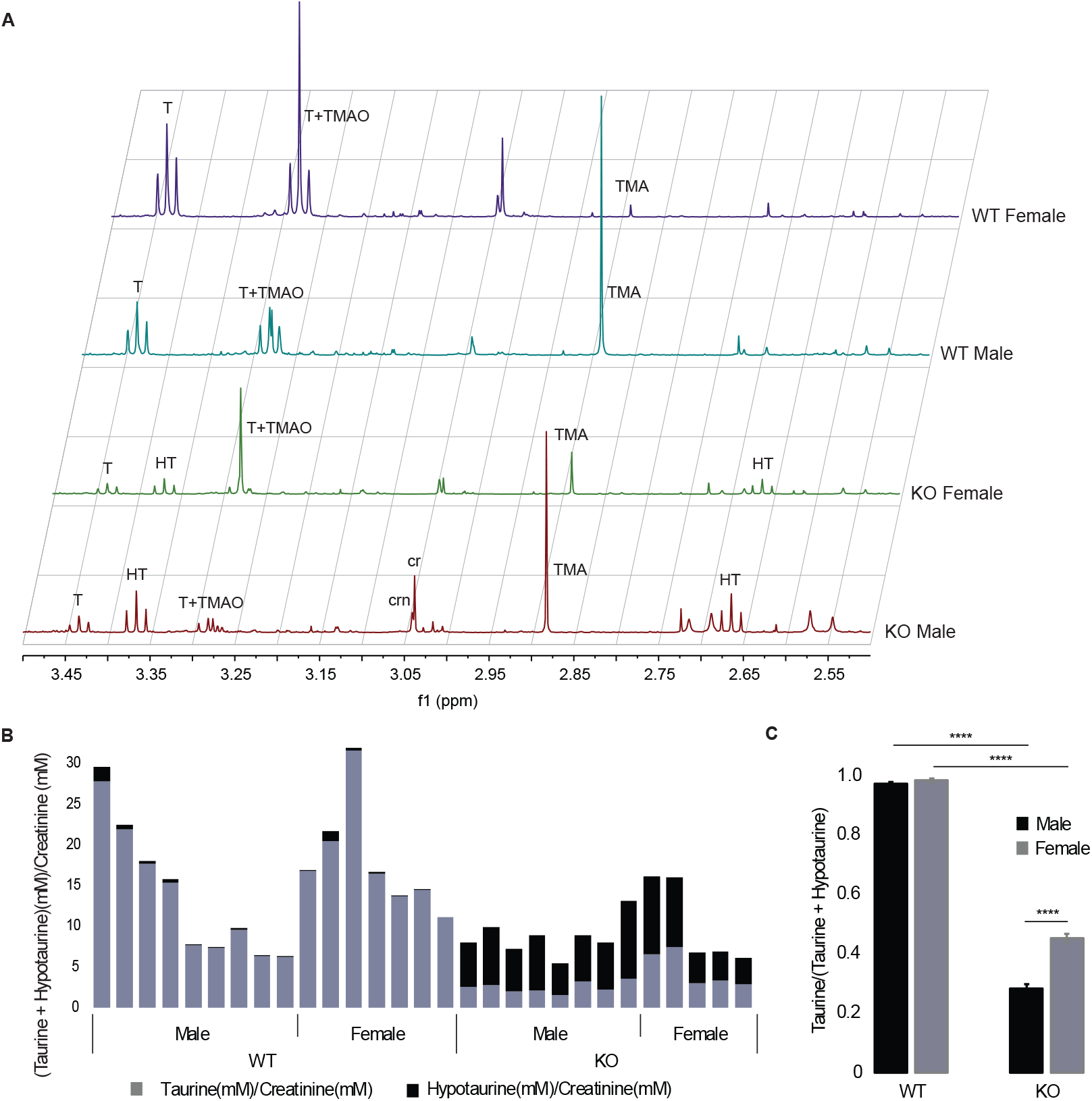
Abundance of hypotaurine and taurine in mouse urine. (A) Representative NMR spectra of urine from *Fmo1*^*−/−*^, *2*^*−/−*^, *4*^*−/−*^ (KO) male and female mice and wild-type (WT) male and female mice. T, taurine; HT, hypotaurine; TMAO, trimethylamine *N*-oxide; TMA, trimethylamine; Crn, creatinine; Cr, creatine. (B) Proportion of taurine and hypotaurine (normalized to creatinine) in urine from individual male and female WT and KO mice. (C) Average ratios of taurine to taurine + hypotaurine in urine of male and female WT and KO mice. Data are expressed as means ± SEM (n = 7-9, WT; 5-8, KO). ****, *P* <0.0001.

Of the three genes deleted in the knockout mouse, the gene encoding FMO4 is expressed at very low levels in mouse (14) and human (15), and that encoding FMO2 is expressed in low amounts in mouse (14) and, in most humans, the gene does not encode a functional protein because of the presence of a premature stop codon (16). In contrast, the gene encoding FMO1 is relatively highly expressed in a number of tissues in both mouse (14) and human (15). Therefore, in humans the most likely candidate for catalyzing the oxygenation of hypotaurine to produce taurine is FMO1. To investigate whether this was the case, baculosomes containing recombinantly expressed human FMO1 were incubated with hypotaurine and the cofactor NADPH at pH 8.5, the optimum for FMO1, and at the more physiological pH of 7.4. Analysis of reaction products by 1D ^1^H NMR spectroscopy identified signals at 3.276 and 3.433 ppm, corresponding to taurine, at both pHs (Fig. 2A). As expected of an FMO1-catalyzed reaction, with NADPH as cofactor, production of taurine was greater at pH 8.5 than at pH 7.4 (*P* <0.01) (Fig. 2B). In comparison, very little taurine was detected in incubations of control insect cell microsomes at pH 8.5 and none at pH 7.4 (Fig. 2A, B). The identity of taurine as a product of FMO1-catalyzed reactions was confirmed by high-resolution UPLC-electrospray mass spectrometry against an authentic reference standard (Fig. S2) (17).

**Figure 2.**
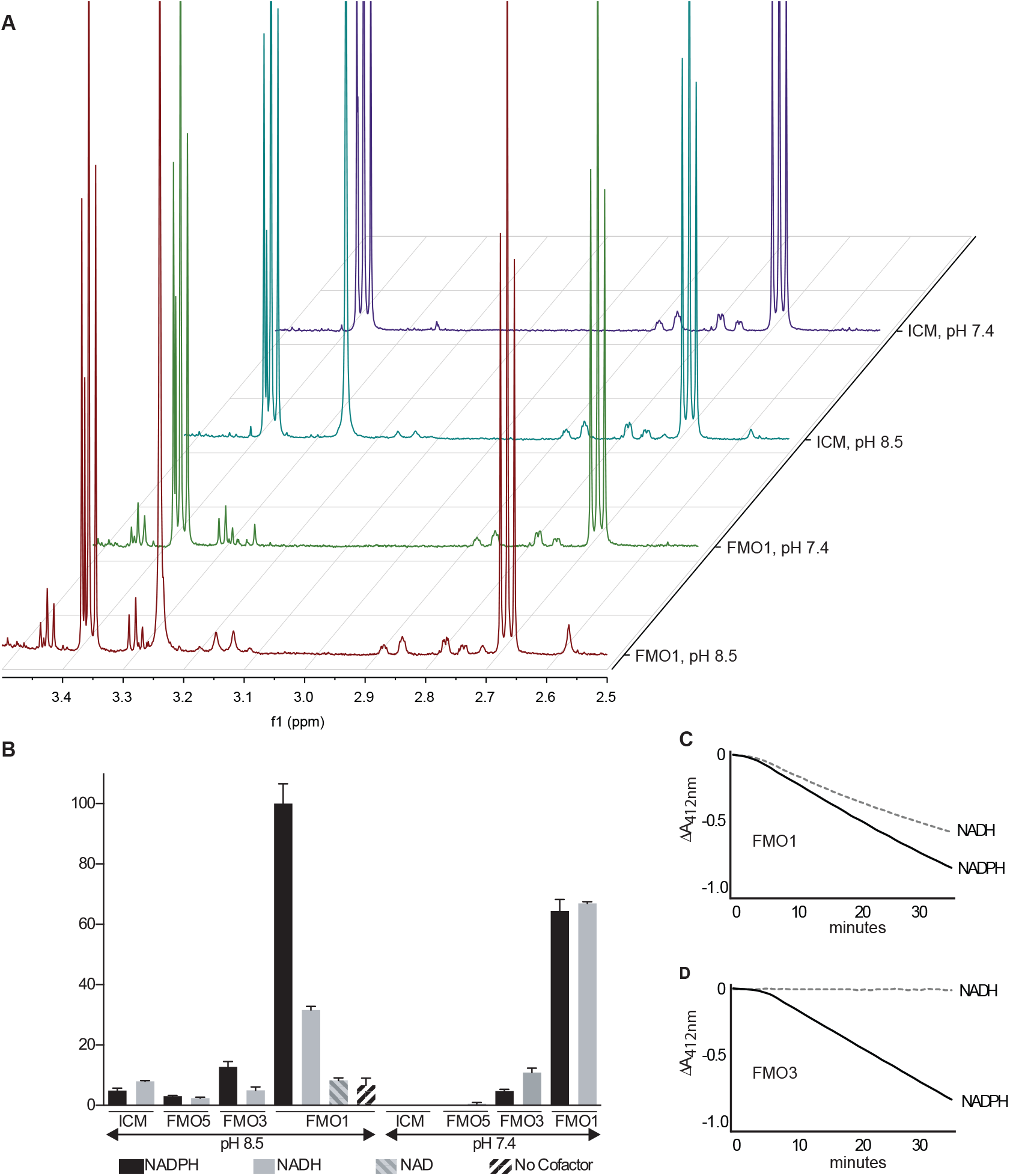
Analysis of in vitro enzyme-catalyzed reactions. (A) Representative NMR spectra of products of reactions catalyzed by baculosomes containing human FMO1 or by insect cell microsomes (ICM) at pH 8.5 or 7.4. Note: the singlet signal at ca 3.364 ppm overlapping the triplet due to hypotaurine is due to methanol. T, taurine; HT, hypotaurine. (B) Production of taurine from hypotaurine in reactions catalyzed by human FMO1, FMO3 or FMO5 or by ICM, at pH 8.5 or 7.4, with NADPH or NADH as cofactor. In the case of reactions catalyzed by FMO1 at pH 8.5, NAD or no cofactor were also used. Taurine production was quantified by NMR and is plotted relative to that produced by FMO1-containing baculosomes at pH 8.5 with NADPH as cofactor (set at 100%). Data are expressed as means ± 0.5 × range (n =2). (C, D) Progress curves of methimazole *S*-oxygenation catalyzed by human FMO1 (C) or FMO3 (D), at pH 8.5, with NADPH or NADH as cofactor. Reactions were monitored at A_412nm_.

We also investigated whether the conversion of hypotaurine to taurine could be catalyzed by either of the two other major functional FMOs of humans, FMO3 and FMO5 (18). In comparison with baculosomes containing human FMO1, those containing FMO3 produced much lower amounts of taurine at pH 8.5 or 7.4, with either NADPH or NADH as cofactor (*P* <0.01). Production of taurine by baculosomes containing human FMO5 was significantly less than by FMO3-baculosomes (*P* <0.05) and was similar to that produced by control insect cell microsomes (Fig. 2B). These results confirm that production of taurine from hypotaurine can be catalyzed by FMO1 and, to a much smaller extent, by FMO3, but not significantly by FMO5.

The production of taurine in the *Fmo1*-null mice was less affected in females (~55% depletion) than in males (~70% depletion) (*P* <0.0001) (Fig. 1C). This gender difference is likely due to the contribution in female *Fmo1*-null mice of FMO3, which is absent from the liver of adult male mice (14,19). In the case of wild-type mice, the presence of FMO1, which is more effective than FMO3 in catalyzing the formation of taurine from hypotaurine, as evidenced by analysis *in vitro* (Fig. 2B), greatly outweighs the effect of the presence in females of FMO3.

FMOs, despite being termed NADPH-dependent monooxygenases, have been reported to be able also to use NADH as a cofactor (20). We found that in catalyzing the production of taurine from hypotaurine, FMO1 could use either NADPH or NADH as cofactor (Fig. 2B). However, whereas NADPH is the more effective cofactor at pH 8.5 (*P* <0.01), at pH 7.4 the cofactors are equally effective (Fig. 2B). When NAD was used as cofactor for FMO1 the amount of taurine produced was very low and not significantly different from that produced in the absence of cofactor (Fig. 2B). FMO1 could also use NADH as cofactor for *S*-oxygenation of the FMO model substrate methimazole (Fig. 2C). However, in the case of FMO3, methimazole oxygenation was dependent on NADPH (Fig. 2D).

Kinetic parameters of FMO1-catalyzed oxygenation of hypotaurine to produce taurine were estimated under conditions at which the enzyme was most active: pH 8.5 with NADPH as cofactor (Fig. 2B). ^1^H NMR spectroscopic analysis revealed that under these conditions there was a 1:1 ratio of taurine production to NADPH oxidation. The kinetics of the enzyme-catalyzed reaction could therefore be assessed by monitoring depletion of NADPH (measured at A_340nm_). A plot of kinetic data is shown in Fig. 3A. Direct linear plots of data (velocity versus substrate concentration) in parameter space gave estimates for *K*_M_ of ~ 4.1 mM and V_max_ of ~ 7.5 μM min^−1^, giving a *k*_cat_ of ~ 55 min^−1^. Further support for the ability of FMO1 to utilize hypotaurine as a substrate is provided by the finding that hypotaurine, at concentrations comparable to the *K*_M_ of FMO1-catalyzed hypotaurine oxygenation, acted as an effective competitor of FMO1-catalyzed *S*-oxygenation of methimazole (Fig. 3B).

**Figure. 3.**
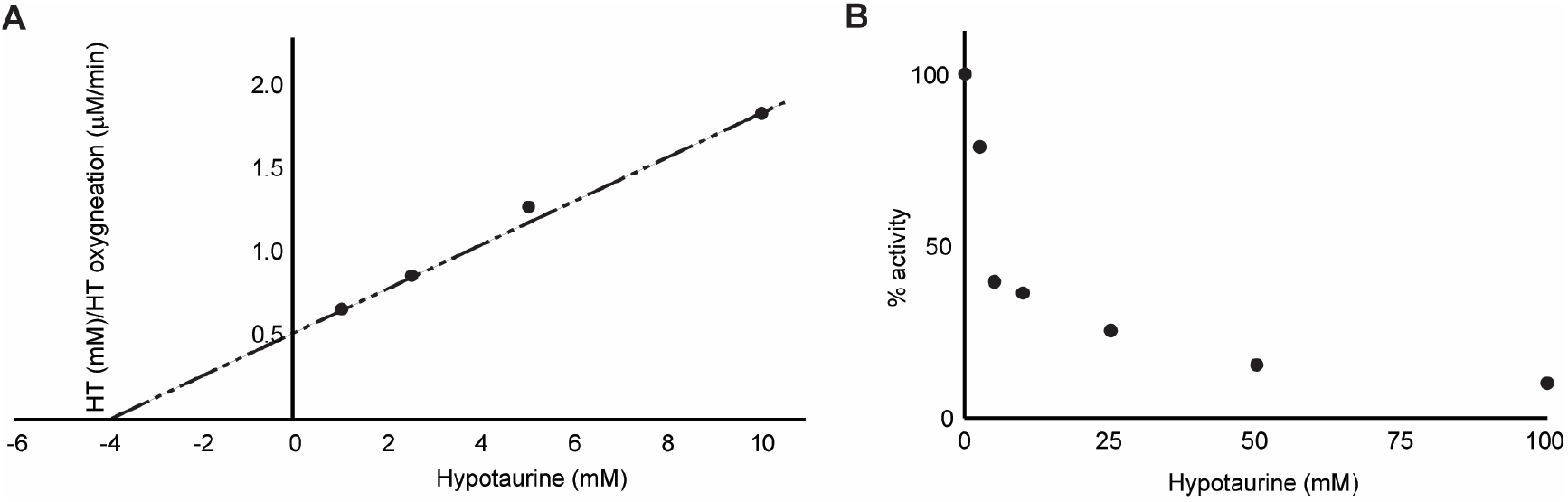
Kinetic and competition analysis of FMO1-catalyzed reactions. (A) Kinetic analysis of FMO1-catalyzed oxygenation of hypotaurine. Reaction mixtures contained human FMO1, NADPH and various concentrations of hypotaurine. Initial velocity was measured by monitoring hypotaurine-dependent depletion of NADPH at A_340nm_. (B) Competition of FMO1-catalyzed *S*-oxygenation of methimazole by hypotaurine. Reaction mixtures contained human FMO1, methimazole, NADPH and various concentrations of hypotaurine. The concentration of methimazole was 50% of the *K*_M_ of FMO1 for this substrate. Initial velocity was measured as described in Experimental Procedures. Methimazole *S*-oxygenation activity was plotted as a percentage of that in the absence of hypotaurine.

## Discussion

We have confirmed both *in vivo*, by ^1^H NMR metabolite profiling of the urine of *Fmo1*^*−/−*^, *2*^*−/−*^, *4*^*−/−*^ mice, and *in vitro*, by analysis of the catalytic activity of FMOs of humans, that formation of taurine from hypotaurine is catalyzed by FMO1, a monooxygenase, and that the enzyme can utilize either NADPH or NADH as cofactor. Our results from knockout mice show that in the absence of FMO1 most taurine production is abolished in both males and females, suggesting that the major source of this abundant amino acid is the FMO1-catalyzed oxygenation of hypotaurine (Fig. 4). Consistent with this, the lack of production of taurine from hypotaurine *in vitro* in the absence of enzyme indicates that non-enzymatic conversion does not contribute substantially to taurine production.

**Figure 4.**
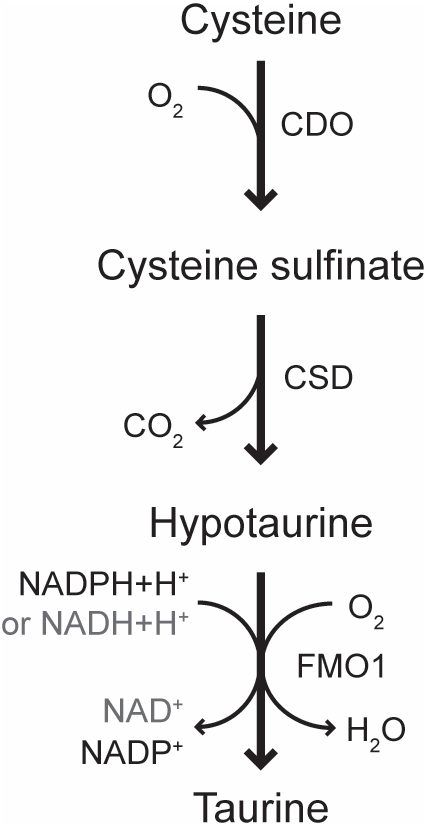
Terminal steps of the biosynthetic pathway of taurine from cysteine. The final reaction in the pathway is catalyzed by FMO1, using either NADPH or NADH as cofactor. CDO, cysteine dioxygenase; CSD, cysteine sulfinate decarboxylase; FMO1, flavin-containing monooxygenase 1.

FMOs (EC 1.14.13.8) are best known for their role in the metabolism of xenobiotics, including therapeutic drugs (reviewed in (21)) and foreign chemicals such as organophosphate insecticides (reviewed in (22)). Of the members of the FMO family, FMO1 has the broadest substrate range (reviewed in (22)). In addition to its role in xenobiotic metabolism, FMO1 has been identified as a novel regulator of energy balance (12).

In humans, *FMO1* is expressed in a range of tissues in which the action of taurine has been implicated, for example, kidney (15), brain (23), small intestine (24), heart (25) and a number of endocrine tissues, including pancreas, adrenal and testis (23). In human liver, *FMO1* is expressed in the fetus, but is switched off after birth (15,26,27). This pattern of expression is consistent with the decline in taurine concentration in the liver after birth (28). In contrast, in adult rodent liver, the gene encoding FMO1 is highly expressed (14) and taurine is abundant (9).

Concentrations of taurine are high in human and monkey fetal brain, but fall during development (28). The decline in taurine concentration in developing brain is consistent with the decrease in expression of *Fmo1* in mouse brain during development (14). FMO1 is active in mouse brain, as evidenced by its catalysis of the *N*-oxygenation of the tricyclic antidepressant imipramine (10). FMO1 would therefore be expected to contribute to the production in brain of taurine from the precursor hypotaurine.

Taurine deficiency is implicated in a number of pathological conditions, including cardiomyopathy, muscular abnormalities and renal dysfunction (2). Conversely, taurine supplementation has been reported to have positive effects on health, for instance, in lowering total plasma cholesterol (29) and in overcoming insulin resistance (2). Given our finding that FMO1 catalyzes the formation of taurine from hypotaurine it is of interest that *Fmo1*-null mice exhibit some characteristics in common with those of taurine deficiency: raised plasma concentrations of cholesterol (13) and glucose (12).

The *K*_M_ of FMO1 for hypotaurine is similar to that of cysteine dioxygenase of human, an upstream enzyme in the taurine biosynthetic pathway (Fig. 4), for its substrate cysteine (30). Although the *K*_M_ for the FMO1-catalyzed oxygenation of hypotaurine is high, our results from the knockout-mouse line indicate that FMO1 is physiologically relevant for the production of taurine.

In addition to *Fmo1*, two other genes are deleted in the knockout mouse line, *Fmo2* and *Fmo4*. Most humans are homozygous for a nonsense mutation of *FMO2*, c.1414C>T[p.(Gln472*)], the *FMO2*\**2A* allele, and do not express functional FMO2 (16,31), and in the case of *FMO4* the gene is expressed at very low levels (15). Thus, although we cannot eliminate the possibility that FMO2 or FMO4 are able to catalyze hypotaurine oxygenation *in vitro*, neither of these enzymes is likely to contribute substantially to the production of taurine in humans *in vivo*.

Commensurate with a role for FMO1 in endogenous metabolism, the gene contains few non-synonymous polymorphisms (32), each of which is present at very low frequency (33) and only one has a significant, but substrate-dependent, effect on catalytic activity (34). However, inter-individual variation of up to 5-fold in the expression of *FMO1* in adult human tissues such as kidney (35) and small intestine (24) could affect taurine production and thus contribute to an intracellular deficiency of the amino acid. In addition, the involvement of FMO1 in the metabolism of drugs implies that drug substrates of the enzyme would compete with hypotaurine for available enzyme and thus compromise taurine production, leading to potentially adverse effects on general health and therapeutic response.

### Experimental procedures

#### Animals

All mice used in this study were bred at University College London. The *Fmo1*^*−/−*^, *2*^*−/−*^, *4*^*−/−*^ mouse line was constructed as described previously (10,36). Mice were genotyped as described (37). Mice were given free access to food (a standard chow diet, Teklad Global 18% ProteinRodent Diet, Harlan Laboratories, Inc., Madison, WI) and water. Animal procedures were carried out in accordance with the UK Animal Scientific Procedures Act and with local ethics committee approval (Animal Welfare and Ethical Review Body) and appropriate Home Office Licenses. Urine was collected between 10:00 AM and 12:00 PM (noon) from male and female mice aged 15-16 weeks. Urine samples were immediately frozen on solid CO_2_ and stored at −80 °C until analyzed by ^1^H NMR spectroscopy, as described below.

#### Enzyme assays

All reaction mixtures (final volumes of 250 μl in a Corning Costar 96-well cell-culture plate, VWR, Lutterworth, Leicestershire, UK) were incubated in a Sunrise absorbance microplate reader (Tecan, Grödig, Austria) equipped with Magellan software, v. 6.2. Reaction mixtures contained either 0.1 M potassium phosphate buffer, pH 7.4, or 0.1 M Tris-HCl, pH 8.5, 1 mM EDTA (aerated immediately before use by shaking for 5 min at room temperature), 2.5 mM hypotaurine, 0.5 mM NADPH, NADH or NAD, or no cofactor, and baculosomes containing human FMO1, FMO3 or FMO5 (135 nM final concentration) (Sigma Aldrich, Gillingham, Dorset, UK) or an equivalent amount of control insect cell microsomes (Corning Life Sciences, Woburn, MA). The mixtures were incubated at 37 °C for 60 min. Monitoring of NADPH or NADH depletion, measured at A_340nm_, revealed that reaction velocities were linear over the 60-min period. Samples were stored at −80 °C until analysed by ^1^H NMR spectroscopy, as described below.

Methimazole *S*-oxygenation was measured by the method of Dixit and Roche (38). Reaction mixtures contained final concentrations of 67 nM of human FMO1 or FMO3 in baculosomes (Sigma Aldrich), 2.5 mM methimazole and 0.5 mM NADPH or NADH, and were incubated at 37 °C.

For estimation of kinetic parameters, reaction mixtures were assembled, by adding, to final concentrations, in the following order 0.1 M Tris-HCl (pH 8.5), 1 mM EDTA (aerated immediately before use by shaking at 37 °C for 10 min), 0.5 mM NADPH and baculosomes containing human FMO1 (135 nM final concentration) (Sigma Aldrich). The mixtures were equilibrated at 37 °C for 3 min, to allow formation of the active C4a-peroxyflavin species of the FAD prosthetic group of FMO. Reactions were initiated by addition of hypotaurine, to final concentrations of 1 to 10 mM or, in the case of blank samples, an equivalent volume of buffer was added. Reaction mixtures were incubated at 37 °C. The initial velocity of enzyme-catalyzed reactions was assessed by monitoring the depletion of NADPH, measured at A_340nm_. Δ A_340nm_ was converted to Δ NADPH using a molar extinction coefficient of 6.2 × 10^3^ M^−1^ cm^−1^ and a light-path length of 0.73 cm. To determine substrate (hypotaurine)-dependent oxygenation of NADPH, readings from a blank sample (a reaction mixture containing no hypotaurine) were subtracted.

The ability of hypotaurine to act as a competitor substrate of FMO1 was assessed by measuring the effect of various concentrations of hypotaurine on FMO1-catalyzed *S*-oxygenation of methimazole. Methimazole *S*-oxygenation was measured by the method of Dixit and Roche (38), as described above, in reaction mixtures containing final concentrations of 67 nM human FMO1 in baculosomes (Sigma Aldrich), 4 μM methimazole and 2.5 to 100 mM hypotaurine.

#### Sample preparation for NMR spectroscopy

Urine samples (50 μl) were prepared for NMR spectroscopy as described previously (39). Enzyme assay samples were thawed, vortexed, then 160 μl of sample was mixed with 80 μl of 0.6 M phosphate buffer, as described previously (39). The samples were re-vortexed and centrifuged at 13 000 g for 5 min at 4 °C. Supernatant (200 μl) was then pipetted into 3.0-mm outer diameter (o.d.) SampleJet NMR tubes (Norell, S-3.0-500-1).

#### NMR spectroscopic analysis

^1^H NMR spectra of urine and enzyme assay samples were recorded on a Bruker Avance III spectrometer (Bruker BioSpin GmbH, Rheinstetten, Germany) operating at 600.44 MHz and at a temperature of 300.0 K, using a standard 1D NOESY presaturation pulse sequence with gradient pulses (noesygppr1d), as described previously (39).

NMR spectral processing was carried out in MNova (MestReNova, version 12.0.1-20560, Mestrelab Research S.L.). The deconvolution of the peaks for metabolite quantification was done using the MNova GSD algorithm. The peak areas were obtained, and the residuals were manually minimized by adjusting the fitting parameters of each peak. Data were imported into Matlab (R2014b, MathWorks). Statistical analysis was performed using an unpaired two-tailed t-test. Significance level *P* < 0.05.

#### NMR data deposition

Original NMR data will be deposited in MetaboLights (EBI UK) (40) after publication.

#### Metabolite identification

NMR-based metabolite identification was carried out using standard methods, as described (17), and using information from the literature and public databases including the Human Metabolite Database (41) (HMDB, http://www.hmdb.ca/).2018). Hypotaurine in *Fmo1*^*−/−*^, *2*^*−/−*^, *4*^*−/−*^ mouse urine showed the following features: 2.665 (t, 6.9 Hz), 58.6 (HSQC) with HMBC to 36.4 and COSY (see Fig. S1) to 3.365 (t, 6.9 Hz), 36.5 (HSQC) and HMBC to 58.5 ppm, in complete agreement with literature values: 2.66 (t, 6.9 Hz), 58.5 and 3.35 (t, 6.9 Hz), 36.2 ppm (HMDB00965, accessed from http://www.hmdb.ca/spectra/nmr_one_d/1626 on 5 February 2019). Taurine in *Fmo1*^*−/−*^, *2*^*−/−*^, *4*^*−/−*^ and wild-type mouse urine showed the following features: 3.283 (t, 6.6 Hz), 50.6 (HSQC) with HMBC to 38.3 and COSY to 3.433 (t, 6.6 Hz), 38.4 (HSQC) and HMBC to 50.5 ppm, in complete agreement with literature values: 3.25 (t, 6.6 Hz), 50.4 and 3.42 (t, 6.6 Hz), 38.3 ppm (HMDB0000251, accessed from http://www.hmdb.ca/spectra/nmr_one_d/1277 on 5th February 2019). Both hypotaurine and taurine in the urine samples were unambiguously identified using the recent MICE criteria (42).^.^

## Supporting information

Veeravalli et al Supplementa Figures

## Acknowledgments

We thank Mr Mohamed Said and Ms Dorna Varshavi for assistance with NMR spectroscopy and Dr Iain Goodall for assistance with UPLC-MS. We also thank Professors Jeremy Nicholson and Elaine Holmes for access to NMR facilities at Imperial College London.

## Author contributions

a complete list of contributions to the paper;

Conceptualization (IRP, JRE, EAS)

Data Curation (SV, IRP, RTF, DV, JRE, EAS,)

Formal analysis (SV, IRP, RTF, JRE, EAS)

Investigation (SV, IRP, EAS, RTF, DV, JRE)

Methodology (IRP, EAS, JRE)

Project administration (IRP, EAS, JRE)

Resources (IRP, EAS, JRE)

Software (RTF, DV)

Supervision (IRP, JRE, EAS)

Validation (SV, IRP, RTF, DV, JRE, EAS)

Visualization (SV, IRP, EAS, RTF, JRE)

Writing - original draft (IRP, EAS)

Writing – review & editing (SV, IRP, RTF, DV, JRE, EAS)

## Data and materials availability

NMR Spectroscopic data will be deposited in MetaboLights (EBI) following acceptance of the manuscript.

