## Supplementary material for "FMO1 catalyzes the production of taurine from hypotaurine": Veeravalli et al Supplementa Figures

**2D NMR spectroscopic analysis**

2D NMR experiments were carried out on selected urine samples to conﬁrm the identification of metabolites (Fig. S1). Representative samples, scaled up to larger volumes, were prepared, as described previously (1), in order to provide greater sensitivity for 2D NMR experiments, including 2D ^1^H JRES (J-resolved), COSY (correlated spectroscopy), TOCSY (total correlation spectroscopy), HSQC (heteronuclear single quantum coherence) and HMBC (heteronuclear multiple bond correlation) NMR. The detailed parameters for these 2D NMR experiments were very similar to those described previously (1).

**Mass spectrometry analysis of products of *in vitro* FMO1-catalyzed reactions**

NMR identification of taurine in the *in vitro* FMO-catalyzed reaction mixtures was additionally confirmed by orthogonal high-resolution ultra-performance liquid chromatography - mass spectrometry (UPLC-MS) analysis (Fig. S2) for unambiguous metabolite identification, according to current Metabolomics Standards Initiative guidelines (2,3). UPLC-MS analysis was carried out using an UPLC system coupled to an accurate-mass Quadrupole Time-of-Flight (Q-TOF) mass spectrometer, as described previously (1), except that the following 7-min gradient program was used, with A = 0.1% aqueous formic acid and B = acetonitrile (ACN) with 0.1% formic acid and A + B = 100% at each timepoint: 5% B at 0 – 4.5 min, 95% B at 4.5 – 5.1 min, followed by re-equilibration from 5.1 to 7.0 min.

Mass spectrometry was performed using a Waters Synapt G2, operating in electrospray ionisation (ESI) mode (negative ion) with lock mass in operation, as described previously (1). Mass detection was carried out in the full-scan mode with an *m/z* range from 50 to 800 in negative-ion mode and a mean resolution of 30,000. Authentic taurine reference standard (Sigma-Aldrich) eluted at 0.50 min and gave an M-H+e- base peak at m/z 124.0071 (mass error 2.4 ppm). Two independent samples of *in vitro* FMO1-catalyzed hypotaurine-containing reaction mixtures gave peaks for taurine at ca 0.50 min, with base peaks at m/z 124.0075 (mass error 0.8 ppm) and 124.0076 (mass error 1.6 ppm), confirming the identity of the reaction products. All output accurate mass measurements were corrected for the mass of the electron, which the commercial software currently fails to do: this is important for low mass metabolites such as taurine.

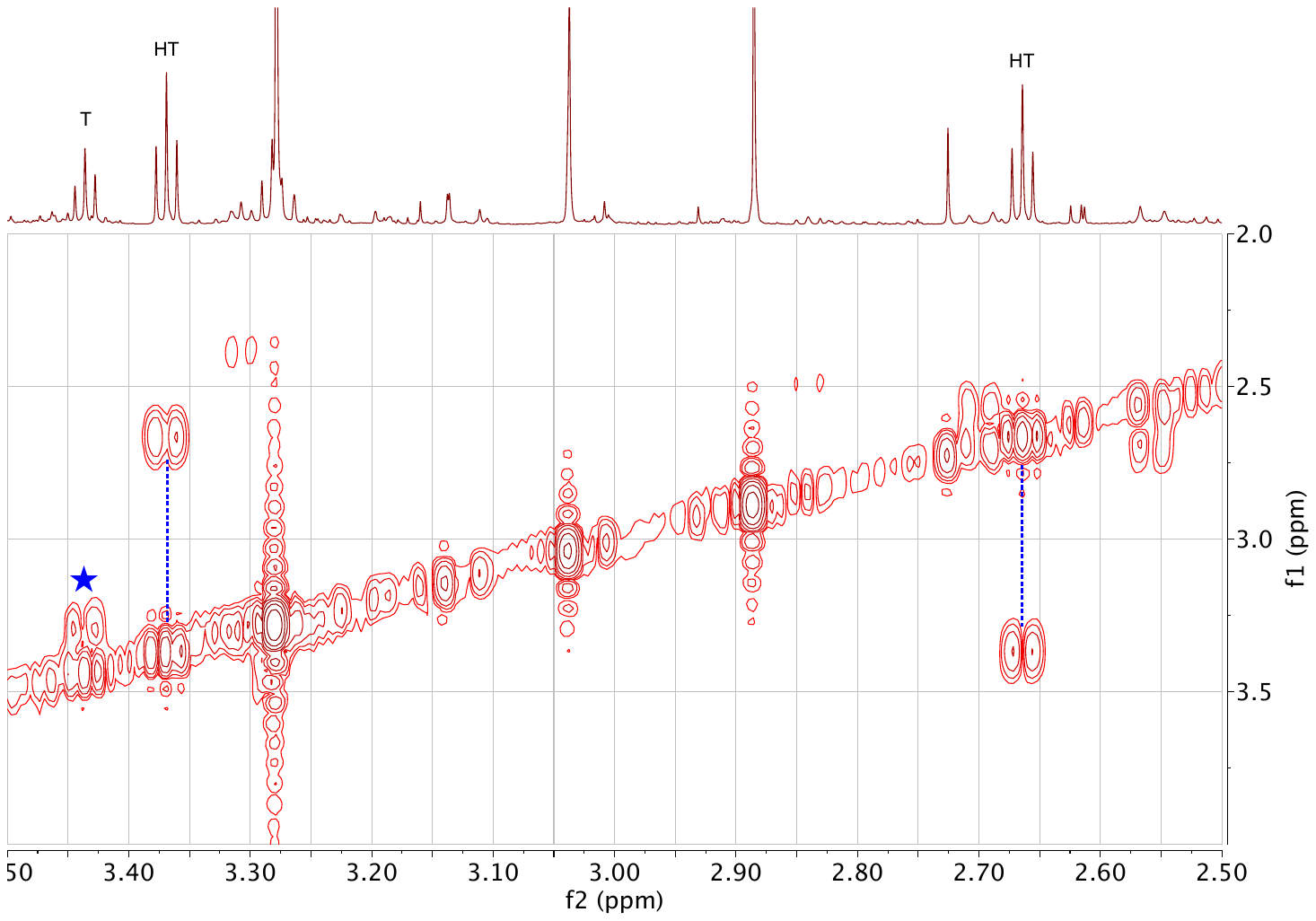

**Figure S1.** An expansion of the aliphatic region of the 600 MHz ^1^H COSY NMR spectrum of the urine from a male, 8-week-old *Fmo1^-/-^, 2^-/-^, 4^-/-^* mouse in the region of the signals for taurine (T) and hypotaurine (HT), plotted as a contour map underneath the corresponding region (3.50 to 2.50 ppm) of the 1D ^1^H NMR spectrum of the same sample. As expected, the two triplet resonances for hypotaurine are connected via off-diagonal COSY cross-peaks indicated by the dashed blue lines connecting the cross-peak to the corresponding f2 signal. A cross-peak connecting the two triplet resonances of taurine (ca 3.43 and ca 3.28 ppm) is indicated by a blue star (★): the other cross-peak is partly hidden by the t1 noise at ca 3.28 ppm.

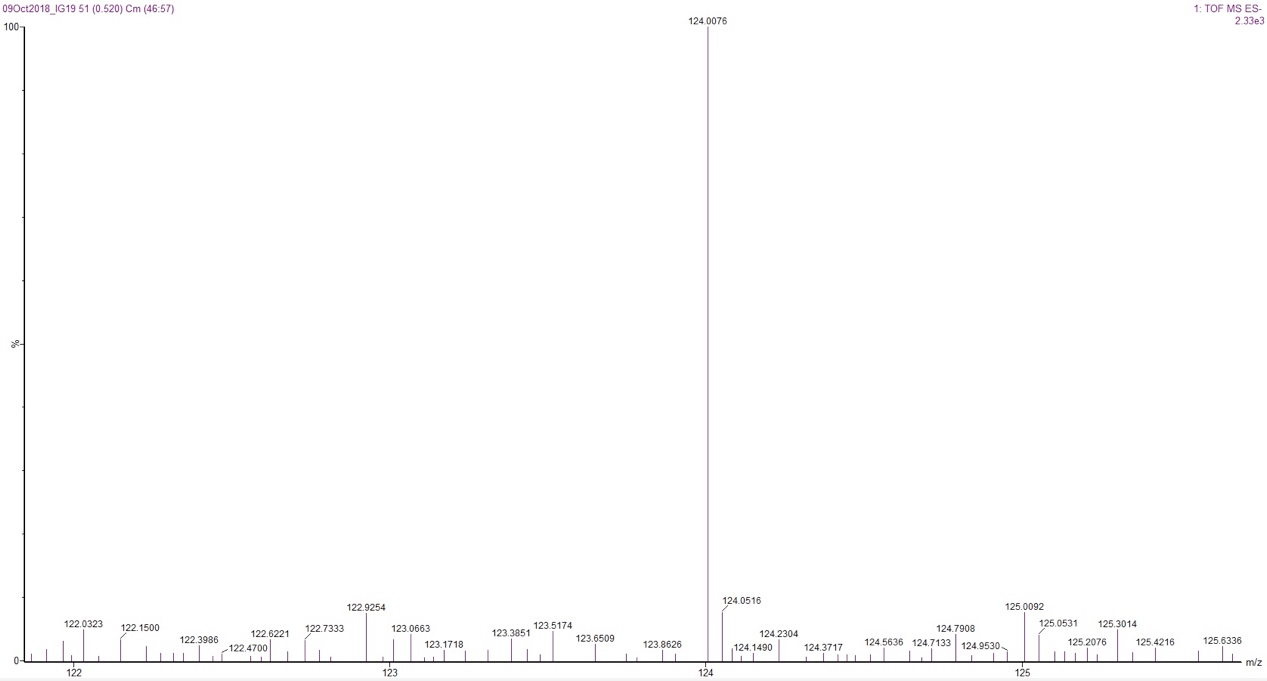

Figure S2. Expansion of the high-resolution negative-ion electrospray mass spectrum of a product peak eluting at ca 0.5 min from an *in vitro* FMO1-catalyzed reaction with hypotaurine as substrate, showing a peak at m/z 124.0076 corresponding to the M-H+e- ion for taurine (expected m/z 124.007385). Authentic taurine eluted also at ca 0.5 min and showed the same spectral features (see Supporting Information text).
